# Forward masking in a bottlenose dolphin *Tursiops truncatus*: Dependence on azimuthal positions of the masker and test sources

**DOI:** 10.1101/2022.01.11.475800

**Authors:** Vladimir V. Popov, Dmitry I. Nechaev, Alexander Ya. Supin, Evgeniya V. Sysueva

## Abstract

Forward masking was investigated by the auditory evoked potentials (AEP) method in a bottlenose dolphin *Tursiops truncatus* using stimulation by two successive acoustic pulses (the masker and test) projected from spatially separated sources. The positions of the two sound sources either coincided with or were symmetrical relative to the head axis at azimuths from 0 to ±90°. AEPs were recorded either from the vertex or from the lateral head surface next to the auditory meatus. In the last case, the test source was ipsilateral to the recording side, whereas the masker source was either ipsi- or contralateral. For lateral recording, AEP release from masking (recovery) was slower for the ipsi-than for the contralateral masker source position. For vertex recording, AEP recovery was equal both for the coinciding positions of the masker and test sources and for their symmetrical positions relative to the head axis. The data indicate that at higher levels of the auditory system of the-dolphin, binaural convergence makes the forward masking nearly equal for ipsi- and contralateral positions of the masker and test.

## Introduction

The unique hearing abilities of odontocetes (toothed whales, dolphins, and porpoises) have been investigated in many respects [1, 2]. Nonetheless, several mechanisms responsible for these unique abilities have not been elucidated, particularly concerning spatial hearing and its relation to temporal analysis in odontocetes.

As a result of behavioral and auditory evoked potential (AEP) investigations, a number of characteristics of spatial hearing determined in odontocetes are known. In particular, the acuity of binaural [3–8] and monaural [9, 10] receiving beams, the spatial discrimination capability in terms of minimum audible angle [11, 12], and the value of interaural intensity difference [9, 10, 13] were determined. These data indicate the capability of odontocetes to acutely localize sound sources.

However, real acoustic scenes are typically characterized not only by a variety of sound sources but also by their various temporal interrelations. Therefore, the ability to extract a certain sound signal from numerous background sounds depends on both the spatial distribution of sound sources and their temporal distribution, as well as on the ability of the auditory system to separate sounds based on their spatial and temporal characteristics.

In odontocetes, the AEP of the auditory system to paired sound pulses with varying delays between them has been investigated in several studies [14–19]. At short delays, the response to the second pulse of a pair is suppressed as a result of forward masking. The studies revealed quick recovery (release from forward masking) of the AEP. At equal levels and frequency bands of the conditioning stimulus (the masker) and the test stimulus, the recovery took from 1 to 5 ms depending on the stimuli level; the recovery time may be longer when the masker is of a higher level than the test [14, 18] or shorter when the masker and test are of different frequency bands [16].

To date, all the data on responses to paired stimuli have been investigated in odontocetes when both stimuli were projected from one and the same source position. Regularly, it was at a position along the head midline. These data did not provide information on the ability to respond to signals after a masker projected from another position. In the present study, we attempt to compensate for this deficiency by studying AEPs to paired stimuli when the azimuthal positions of the first (masker) and the second (test) stimuli were different. To achieve this goal, AEPs were recorded at various combinations of (i) azimuthal masker and test source positions, (ii) masker-to-test intervals, and (iii) AEP recording positions (vertex or lateral). A comparison of vertex and lateral AEP recordings was undertaken to discriminate between monaural and binaural processes in the detection of the masked test signals.

## Materials and Methods

### Subject and Facilities

The study was performed at the Utrish Marine Station of the Russian Academy of Sciences on the Black Sea coast (Krasnodar Province). The subject was an adult female bottlenose dolphin *Tursiops truncatus*, 260 cm body length, provisionally 28 years old. Before being used for experiments, the animal was kept in a commercial dolphinarium for 25 years. After two months of experimentation, she was returned back to the dolphinarium.

At the Utrish Marine Station, the animal was kept in a round plastic tank 6 m in diameter, 1.7 m deep, filled with sea water. During the experiments, the water level in the tank was lowered down to 0.4 m. The animal was laid on a stretcher in such a manner that its dorsal head surface was above the water surface and the rostrum tip was in the center of the tank (Fig 1). The animal was kept in such a position during the experiment, which regularly lasted 2 to 3 h. After the experiment, the animal was released, and the water depth in the pool was restored to its normal level of 1.7 m.

**Fig 1.**
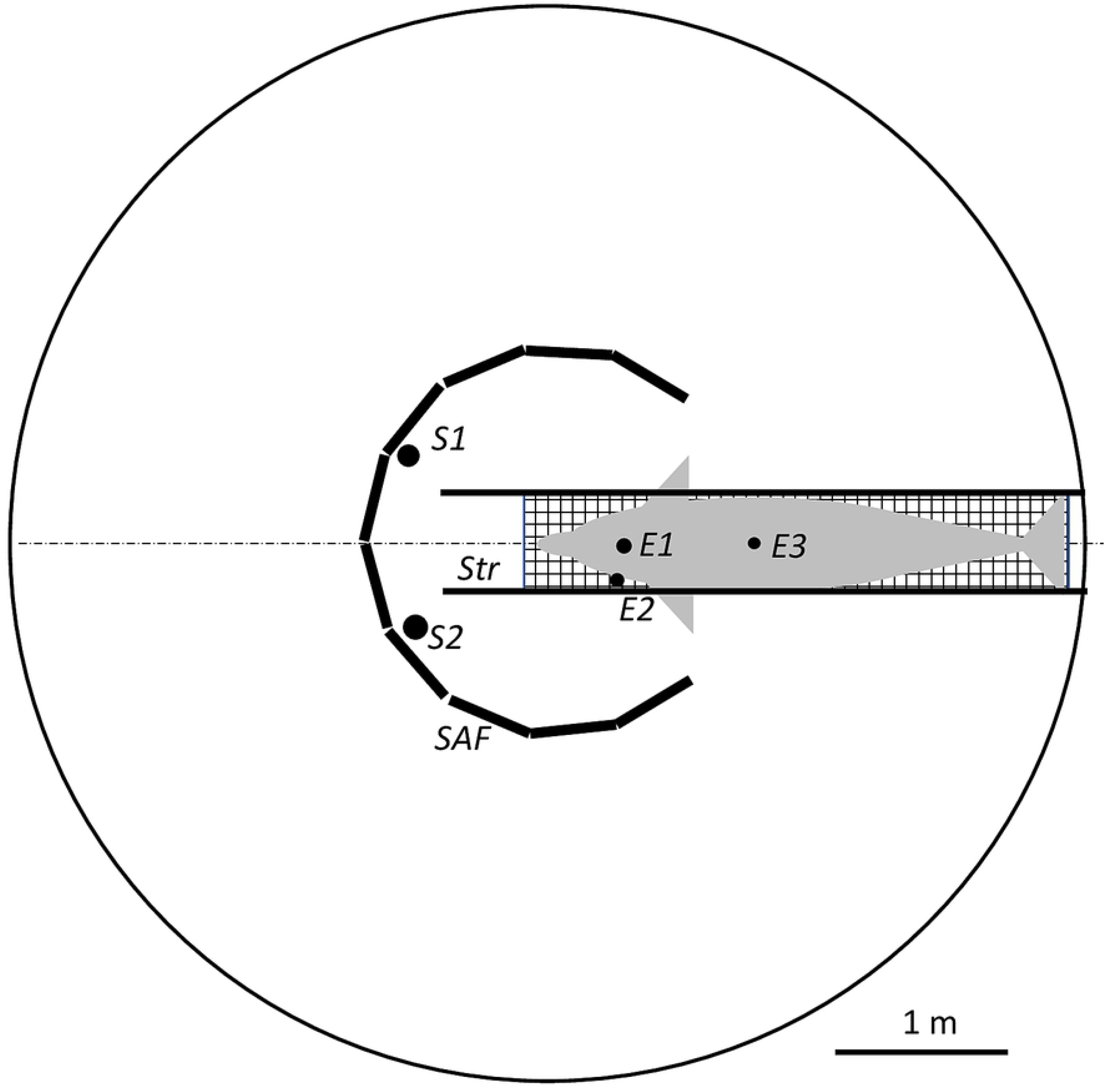
Experimental design. *Str* – stretcher, *E1* – vertex active recording electrode, *E2* – lateral active recording electrode, *E3* – referent electrode, *S1* and *S2* – sound sources, *SAF* – sound absorbing fence.

The study design was approved by The Ethics Committee of the Institute of Ecology and Evolution of the Russian Academy of Sciences (*№* 31, 30.04.2019).

### Acoustic Stimulation

The sound stimuli were tone pips of 64 kHz carriers. The pip envelope was one cycle of a raised cosine function containing 6 cycles of the carrier (Fig 2a, 1). The stimuli were digitally generated by a standard computer at an update rate of 500 kHz using a custom-made program based on the LabVIEW software (National Instruments, Austin, TX, USA). The program generated two pips in separate channels with independent control of the pip amplitudes and interpip delay. The generated signals were digital-to-analog converted by a 16-bit data acquisition board NI USB-6251 (National Instruments), amplified and attenuated by a custom-made two-channel power amplifier-attenuator with a passband of 200 kHz and output impedance of 50 Ohm.

**Fig 2.**
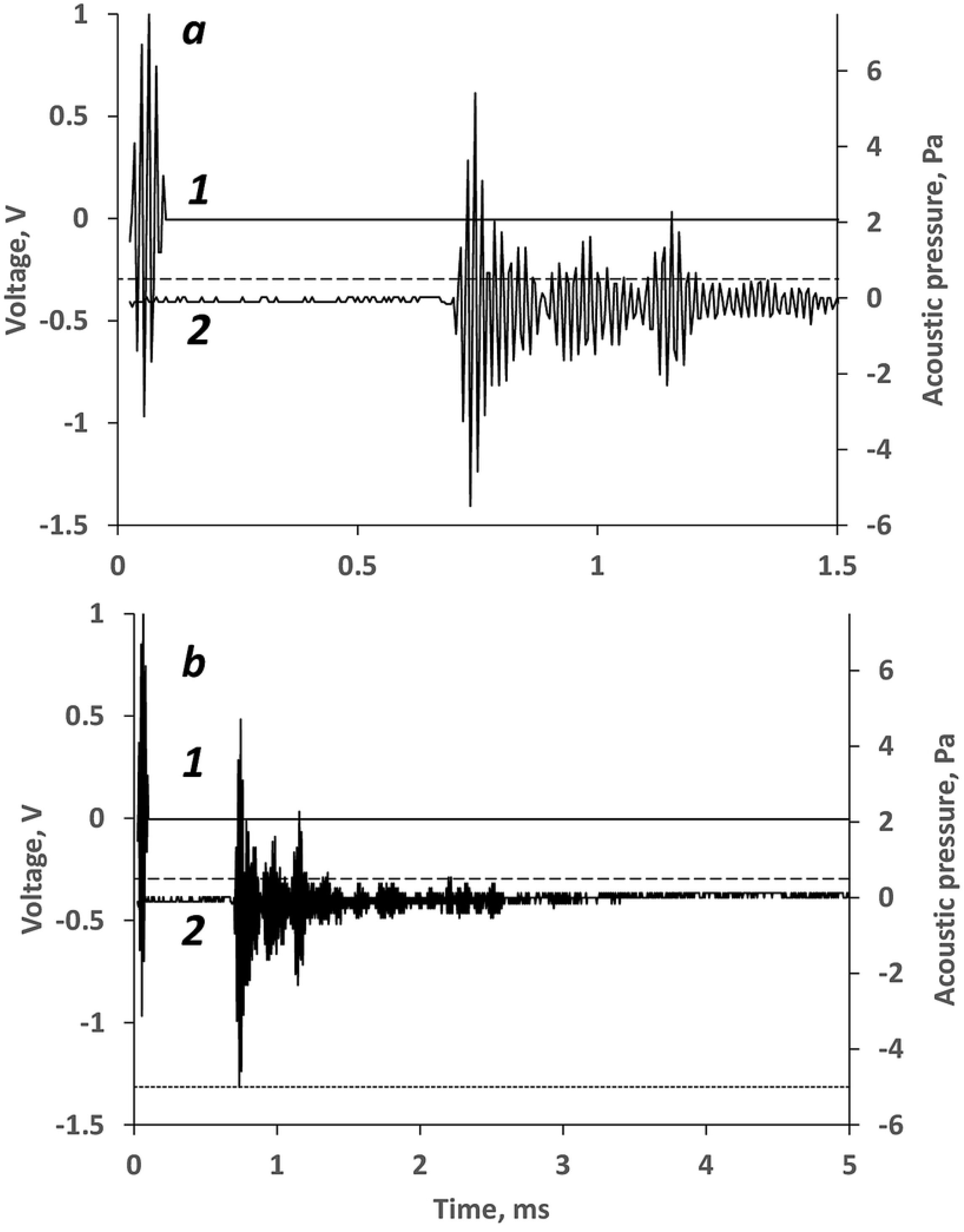
Waveforms of the stimuli. (a) extended time scale, (b) compressed time scale. *1* – electronic signal (left ordinate scale), *2* – acoustic signal recorded from the monitoring hydrophone (right ordinate scale). Dashed lines mark a sound level of –20 dB re. the peak.

The signals generated in two channels fed two B&K 8104 transducers (Bruel & Kjaer, Naerum, Denmark). The transducers were positioned symmetrically with respect to the animal head axis, at a distance of 1 m from the melon tip, at a depth of 20 cm, i.e., at the mid-depth of the water. The azimuthal position of the transducer relative to the head axis varied within a range of ±90°.

To minimize sound reflections from the tank walls, the area around the animal’s head and transducers was surrounded by a circular fence of sound-absorbing material (rubber with closed air cavities) (see Fig 1). The fence covered the whole water depth. The radius of the fence was 1.05 m, so the transducers were positioned inside it.

Stimuli were monitored by a B&K 8103 hydrophone (Bruel & Kjear, Naerum, Denmark) coupled with a Nexus 2690 amplifier (Bruel & Kjaer) and an NI-5132 digital oscilloscope (National Instruments). The monitoring hydrophone was positioned next to the lower jaw of the animal. The acoustic measurements were performed between, not during the AEP-recording sessions.

The waveform of the acoustic signals that were picked up by the monitoring hydrophone did not exactly reproduce the electronic signal that activated the sound-emitting transducers (Fig 2). The acoustic signal was longer than the electronic signal. An initial component of a peak level of 135 dB re. 1 μPa (5.6 Pa acoustic pressure) followed by a few smaller (−6 to −10 dB re. the initial peak) components. During 0.5 ms after the peak, the acoustic pressure fell to approximately −20 dB re. the signal peak.

### AEP Recording

For noninvasive AEP recording, gold-plated electrodes imbedded in silicon suction cups (Cetacean Research Technology, Seattle, WA, USA) were used. Two manners of AEP recording were used: the vertex and lateral.

For vertex AEP recording, one of the electrodes was fastened at the dorsal head surface, 6-9 cm behind the blowhole. This point has been found to feature the highest amplitude of a noninvasively recorded fast AEP known as the auditory brainstem response, ABR [14]. This electrode was considered active for ABR recording. The other electrode was fastened at the dorsal fin where the ABR amplitude was negligible. This electrode was considered reference. Both the electrodes were above the water surface.

For lateral AEP recording, the active electrode was fastened underwater at the lateral head surface, next to the auditory meatus. For this electrode, the embedding suction cup served both as electrode fastening and as insulation from low-impedance sea water. The reference electrode was positioned at the dorsal fin above the water surface.

The electrodes were connected to the input of a brain-potential amplifier LP511 (Grass Technologies) that provided amplification by 80 dB within a frequency range of 0.3 to 3 kHz. This frequency range was chosen as covering the frequency band of ABR but minimally transferring noise outside this band.

Amplified EEG signals were analog-to-digital converted and processed by the same acquisition board NI USB-6251 that was used for signal generation. The sampling rate for conversion was 40 kHz. To process the brain potential signals, a custom-made program based on LabVIEW (National Instrument) software was used.

### Experiment Design and Data Processing

The forward-masking processes were investigated by recording AEPs to two successively presented stimuli, the first of which was considered a masker (conditioning stimulus), and the second was considered a test. The peak levels of both the masker and test stimuli were equal, specifically, 135 dB re. 1 μPa.

To separate the ABRs to the two stimuli, a subtraction procedure was used: the response to the masker alone was subtracted from the response to the pair of stimuli. This procedure requires the responses to the masker to be equal in records obtained with the pairs of stimuli (masker + test) and masker alone. To make this equality as precise as possible, the masker + test pairs and maskers only were presented in an interleaving manner (Fig 3): the masker + test, (stimuli *1* and *2* in Fig 3), the masker only (stimulus *3* in Fig 3), masker + test again, etc. With this manner of presentation, responses to the masker in pairs and maskers only were equally subjected to any long-term variation in hearing sensitivity that might occur from long-term hearing adaptation.

**Fig 3.**
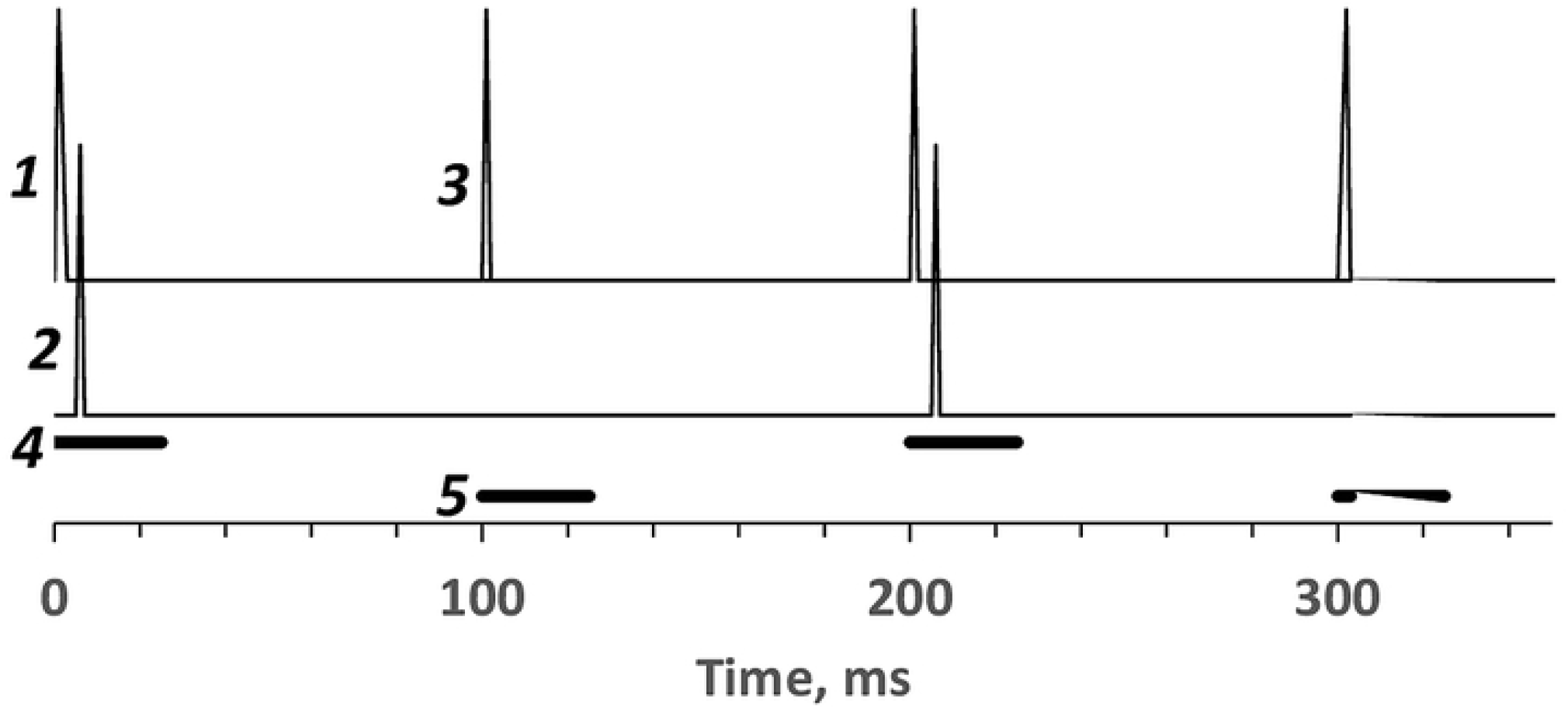
Succession of presentation of masker and test stimuli. A fragment containing two pairs of stimuli and two maskers. The peaks mark presentation of the masker (1) and test (2) in pairs and the masker alone (3). 4 – time windows for collection of AEPs to pairs of stimuli, 5 – time windows for collection of AEPs to the masker alone.

The stimuli were presented at a rate of 10/s. For extraction of AEPs from brain-potential noise, 1000 responses to masker + test pairs and 1000 responses to maskers were collected. Thus, each recording trial lasted 200 s.

AEPs were extracted from brain-potential noise by coherent averaging. AEPs to maskers + test pairs and AEPs to maskers were collected and averaged separately (epochs *4* and *5* in Fig 3, respectively). The averaged AEP to the test stimulus was extracted by point-to-point subtracting the averaged response to the masker from the averaged response to the masker + test. Thus, each recording trial provided three AEP waveforms: (i) the masker, (ii) test, and (iii) masker + test combination.

### AEP amplitude evaluation

The AEP amplitude was assessed as the largest positive-to-negative span within a 2.5-ms window where the response was expected, specifically, from 2.5 to 5 ms after stimulus. A record was considered response-present when this span was 3.5 times as high as the root-mean-square (RMS) of the background record noise. The background noise was evaluated in records obtained using the same procedure as for AEP records but with no stimulus. The criterion of 3.5 RMS was used because for Gaussian noise, RMS is quantitatively equal to the standard deviation (SD) of the amplitude distribution. Therefore, 99.95% of the noise samples are within a range of ±3.5 SD. At a sampling rate of 40 kHz, the 2.5-ms long window contained 100 samples. The probability of background fluctuations exceeding 3.5 SD is 0.9995^100^ = 0.95. Therefore, according to the commonly adopted 95% criterion, voltage spans exceeding 3.5 RMS can be considered not random background noise fluctuation but statistically significant indications of responses.

## Results

### Background noise and AEP waveforms

In both vertex and lateral recordings, the background noise RMS was evaluated as 0.03 μV. Therefore, according to the criterion described above, records were considered response-present if the largest amplitude span within the time window 2.5-5 ms after stimulus exceeded 0.1 μV, and this span was taken as the AEP amplitude.

ABR waveforms consisted of a series of waves of an onset latency (including 0.67-ms acoustic delay) of 2.1 ms and an overall duration of approximately 8 ms (Fig 4). For the vertex recording, the largest positive-to-negative voltage component (2 to 2.5 μV unmasked) had peak latencies of 3.7 ms (negative) and 4.8 ms (positive). Below it is designated as an N3.7-P4.8 component (Fig 4a). With the use of the criterion described above, this component was taken as the AEP amplitude.

**Fig 4.**
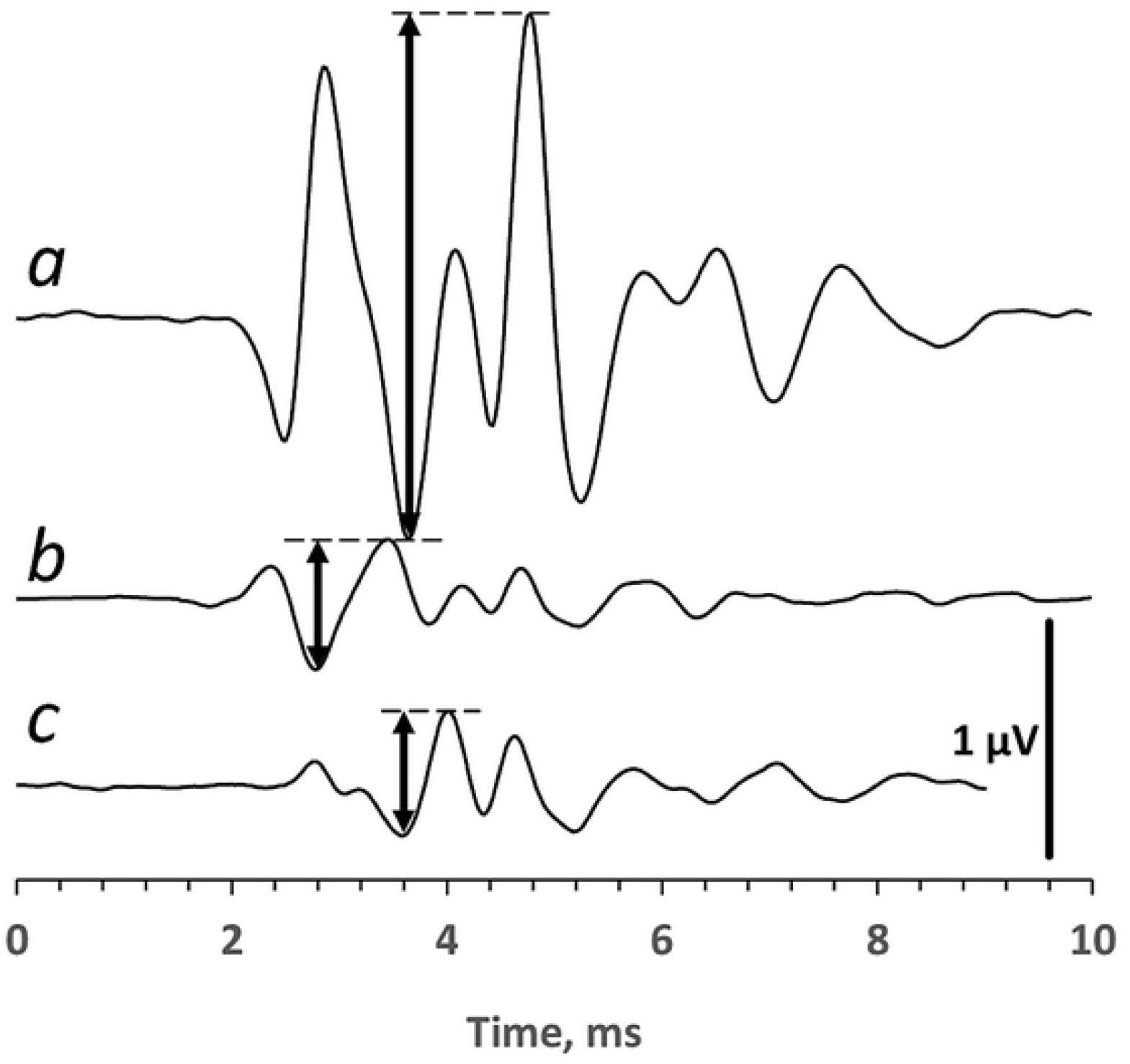
Examples of AEPs recorded from the vertex and lateral head surface. (a) Vertex recording, stimulus source at the head midline. (b) Recording from the lateral head surface with a sound source at 30° ipsilateral to the recording. (c) Recording from the lateral head surface with a sound source at 30° contralateral to the recording. Positive polarity of the active electrode: upward. Double-headed arrows mark the largest negative-to-positive voltages taken as response amplitudes.

For the lateral recording, the AEP waveform depended on the side of sound projection. For ipsilateral stimulation, the earliest positive wave had a peak latency of 2.35 ms (including 0.67-ms acoustic delay). The largest amplitude was characteristic of a negative-positive complex with peak latencies of 2.8 ms (negative) and 3.5 ms (positive) (N2.8-P3.5 component, Fig 4b). Its amplitude was 0.6-0.7 μV unmasked. With the use of the criterion described above, this component was taken as the AEP amplitude.

For contralateral stimulation, the earliest wave had a positive peak latency of 2.8 ms, and the largest amplitude was characteristic of a negative-positive complex with peak latencies of 3.6 ms (negative) and 4.1 ms (positive) (N3.6-P4.1, Fig 4c). With the use of the criterion described above, this component was taken as the AEP amplitude.

### Effectiveness of the subtraction procedure to separate responses to the masker and test

The effect of the subtraction procedure is illustrated in Fig 5 for vertex records obtained with masker-test delays of 1 ms (a) and 5 ms (b). At a shorter delay of 1 ms (Fig 5a), responses to the two stimuli overlapped one another (a, *1*); however, the subtraction procedure extracted a masker-provoked response (a, *2*) and a test response (a, *3*). At a longer delay of 5 ms (Fig 5b), responses to the masker and test did not overlap (record b, *1*); the record b, *2* demonstrates the non-delayed masker response and the absence of a delayed test response, whereas the record b, *3* demonstrates the delayed test response and the absence of a nondelayed masker response.

**Fig 5.**
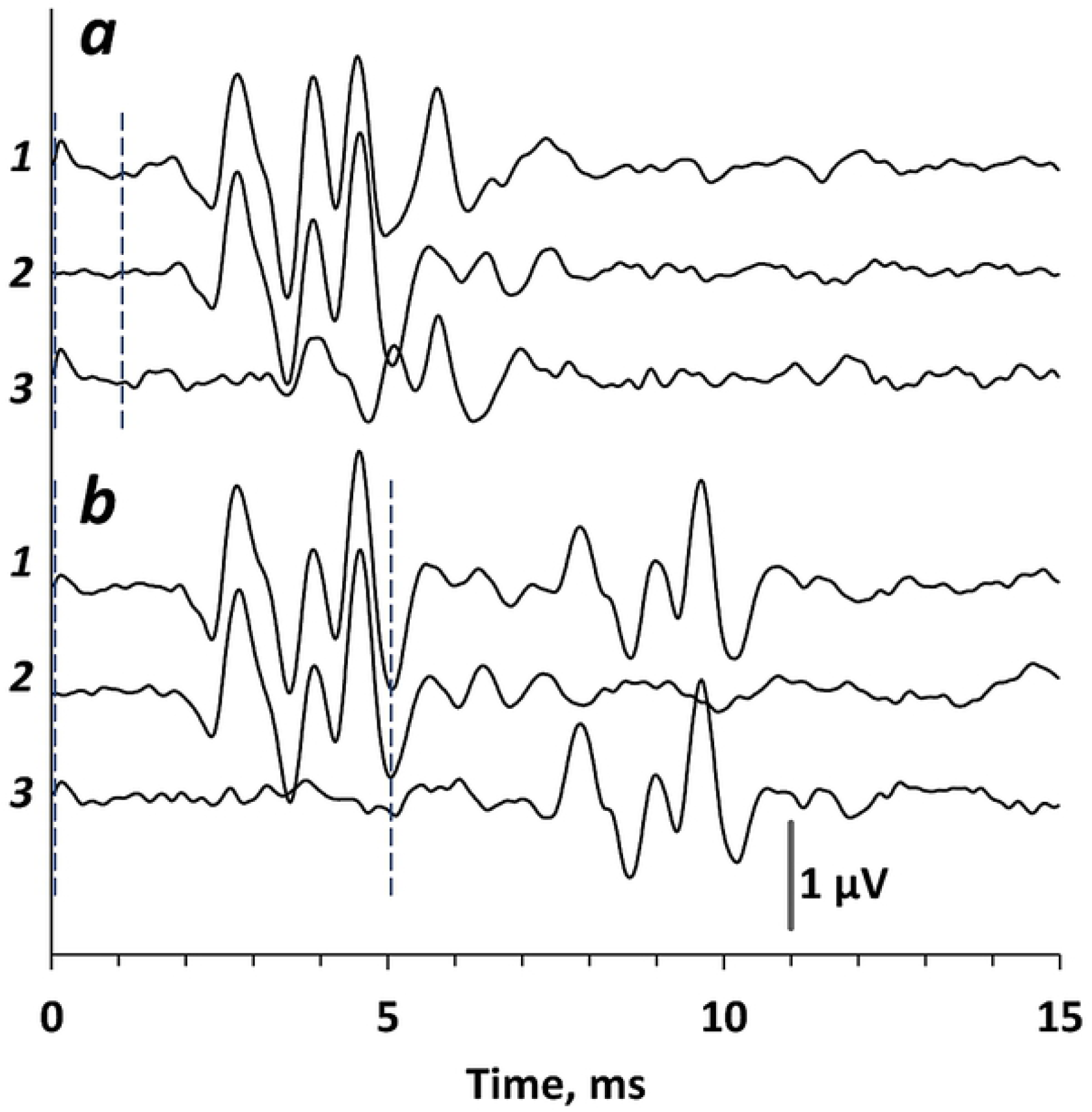
Extraction of the test AEP by the subtraction procedure. The vertex recording and stimulus source are at the head midline. Positive polarity of the active electrode: upward. AEP to a pair of stimuli with 1-ms delay (a) and 5-ms delay (b). The first stimulus (masker) corresponds to the trigger instant, and the instant of the second stimulus (test) is marked by a vertical dashed line. *1* – response to the pair of stimuli. *2* – response to the masker alone. *3* – response to the test extracted by the subtraction of record *2* from record *1*.

### Forward masking of AEP at lateral recordings

For lateral AEP recording, the test stimulus source was positioned ipsilateral to the recording site; the masker source was positioned either ipsilateral or contralateral. Figure 6 exemplifies the records obtained when the masker source was located either at the ipsilateral azimuth of 30° (a) or at the contralateral azimuth of 30° (b). The measurements were repeated three times, and the exemplified records present the average of the obtained original records.

**Fig 6.**
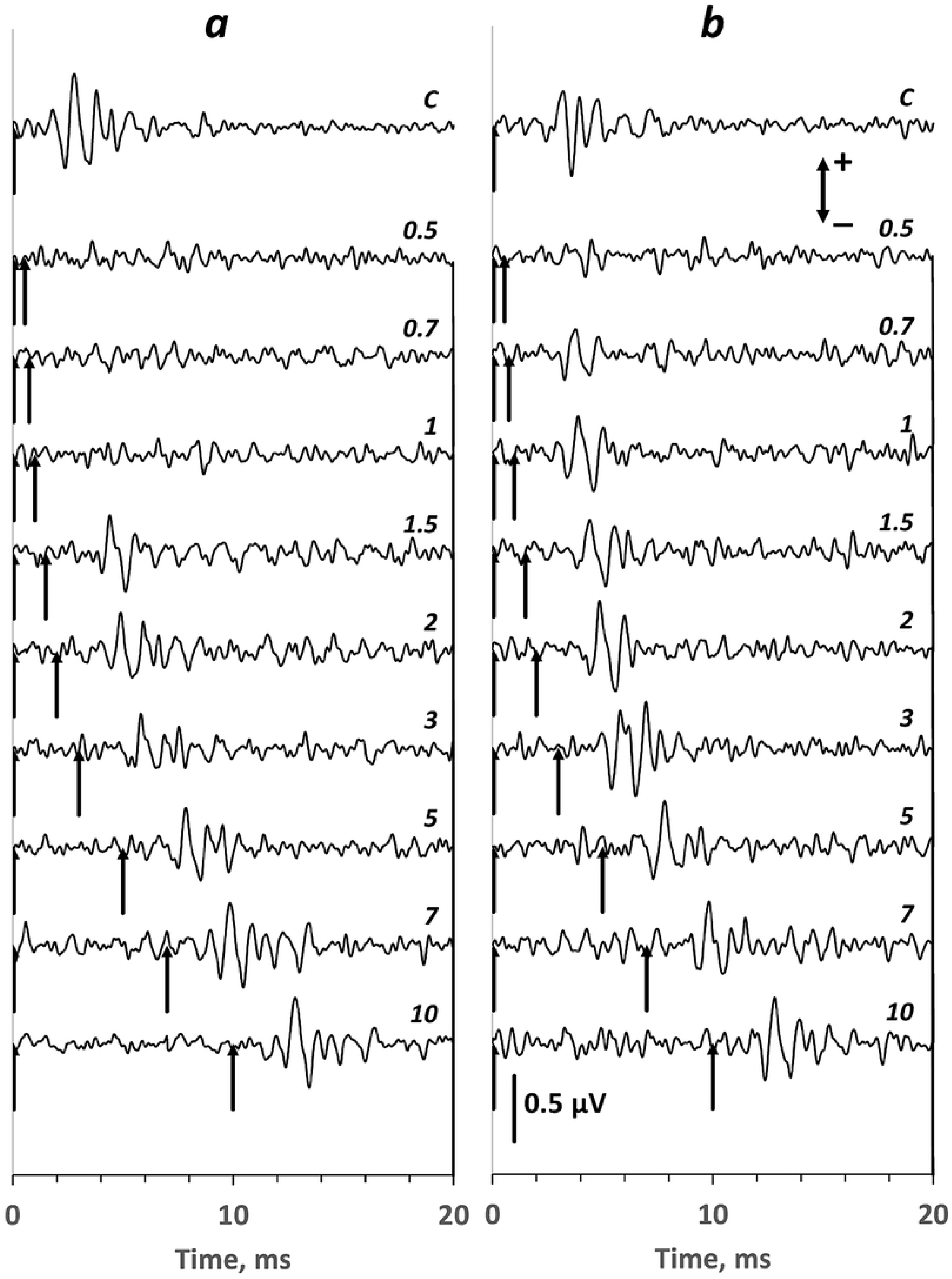
Forward masking effect on laterally recorded AEPs. The records present responses to the second (test) stimulus extracted with the use of the procedure described above. Positive polarity of the active electrode: upward. (a) – masker source ipsilateral to the test. (b) – masker source contralateral to the test. In both cases, the test source was ipsilateral to the recording side. Upward-headed arrows mark stimuli. Interstimulus intervals are shown next to records in ms. (c) – control.

The records illustrate a stronger forward-masking effect for the ipsilateral masker source position than that for the contralateral position. At the ipsilateral masker source position, the main AEP component N2.8-P3.5 appeared at delays no shorter than 1.5 ms. At the contralateral masker source position, this complex appeared at a delay of 0.7 ms; at a delay of 1 ms, its amplitude was close to the control.

All the data obtained with lateral AEP recordings are presented in Fig 7 as dependencies of the test AEP amplitude on the masker-to-test delay. The amplitude measures are those of the largest AEP component, i.e., the N2.8-P3.5 component characteristic of ipsilateral recordings (see Fig 4b). The shortest delay used between the masker and test was 0.5 ms because at shorter delays, the masker “tail” could overlap the test, thus producing simultaneous masking. At a delay of 0.5 ms or longer, the level of the masker was not higher than −20 dB re. the signal peak (see Fig 2); we supposed this level negligible for masking. The longest delay between the masker and test that we used was 7 ms because this delay resulted in complete release of the test from masking.

**Figure 7.**
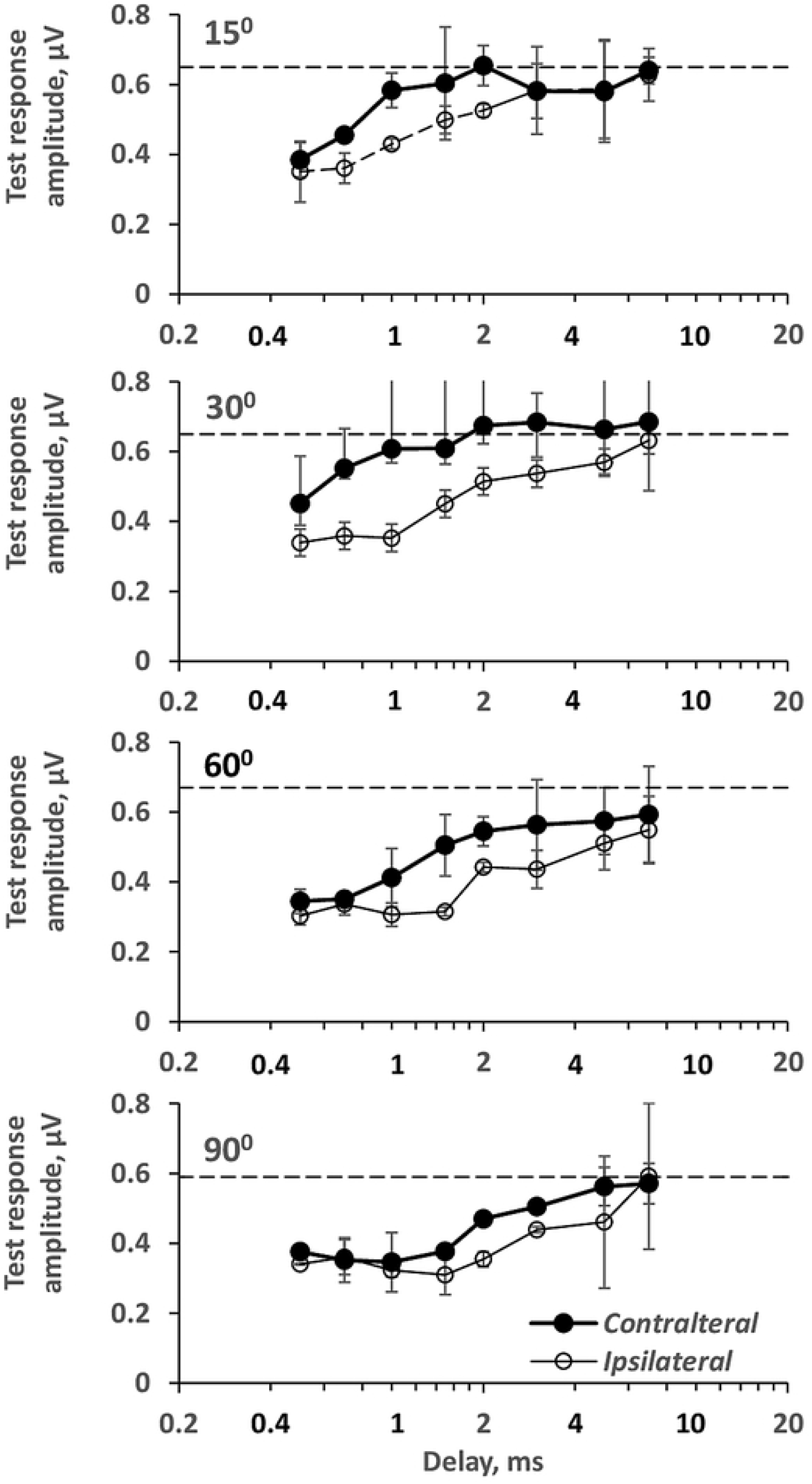
Test AEP amplitude dependence on masker-to-test delay for lateral AEP recordings. Data for azimuths from 15 to 90°, as indicated in the panels. The test stimulus was always ipsilateral to the side of AEP recording. Contralateral: the masker stimulus source was contralateral, symmetrical to the test stimulus source; Ipsilateral: the masker stimulus source was at the same position as the test source. Error bars – interexperiment standard deviations. Dashed lines – control amplitude of AEP to the masker stimulus that had identical parameters to the test stimulus.

For all tested azimuthal positions from 15 to 90°, test AEPs were quicker released from masking at the contralateral than at the ipsilateral masker source position. The differences between recovery functions for contra- and ipsilateral masker source positions were statistically significant (*p* < 0.001) for all tested azimuths from 15 to 90°, as assessed by ANOVA.

### Forward masking of AEP at vertex recordings

Vertex recordings were performed both with the test stimulus source on the right and the test source on the left, and vice versa; records from the two symmetrical test source positions were averaged (Fig 8). Records were obtained when the masker and test sources were located symmetrically at opposite sides (a) and when they were at one and the same azimuth, in close vicinity to one another (b). In the presented case, measurements were performed with a test source positioned at azimuths of +30° and −30°; records were averaged.

**Fig 8.**
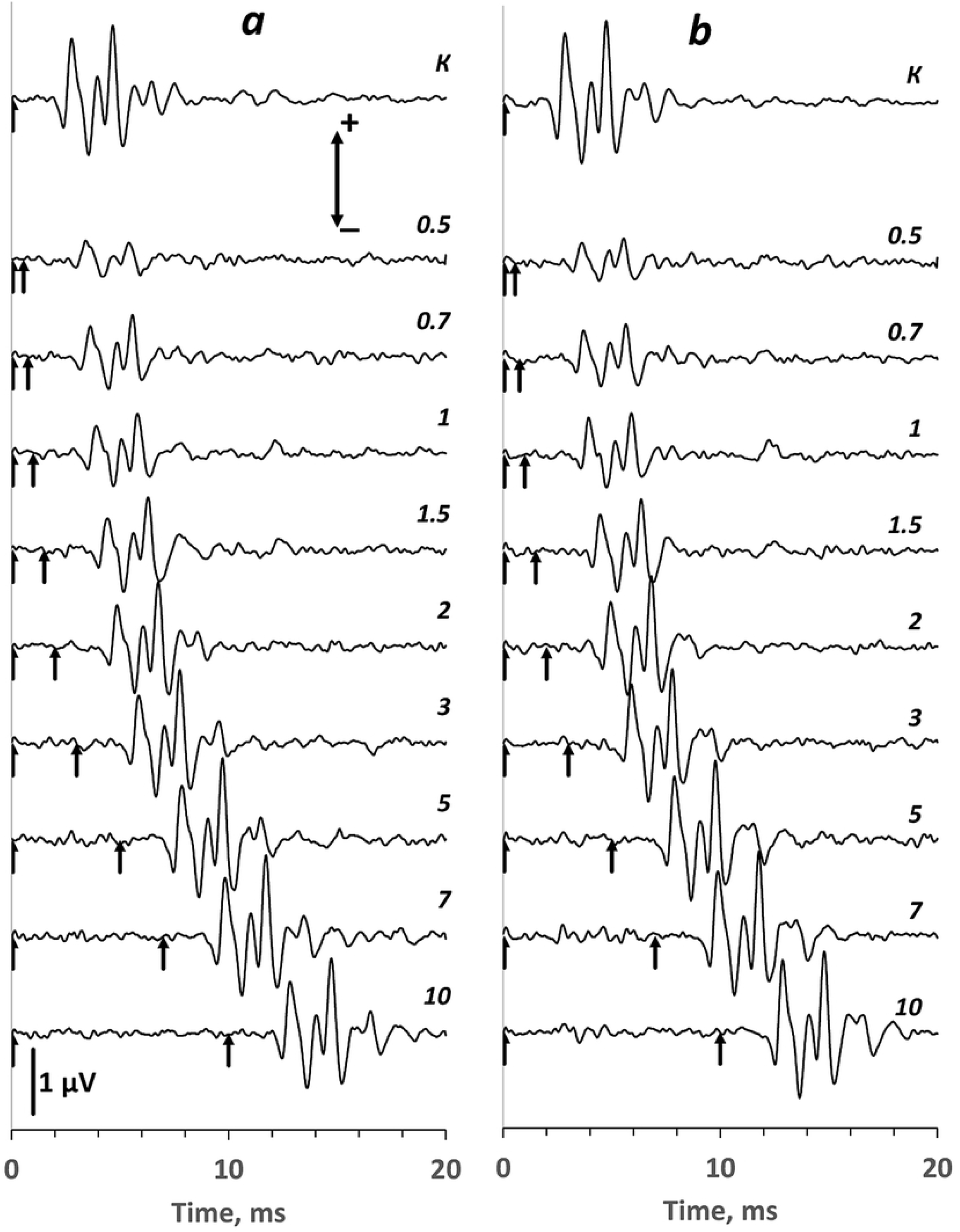
Forward masking effect on vertex-recorded recorded AEPs. The same designation as in Fig 6.

At short delays, the amplitudes of the test (second in the pair) AEPs were decreased compared to that of the control AEP. With increasing delay, the test AEPs recovered (released from the forward masking). The recovery was nearly completed at delays of 3 to 5 ms; at a further increase in the delay, ABR amplitudes were nearly constant and close to the control. This course of recovery was nearly identical both for the masker and test stimuli emitted from opposite sides relative to the head axis (Fig 8a) and for both stimuli emitted from one and the same azimuth (Fig 8b).

All the data obtained with vertex AEP recordings are presented in Fig 9 as dependencies of the test AEP amplitude on the masker-to-test delay. The amplitude measures present those of the N3.7-P4.8 component (see Fig 4a). Similar to the experiments with lateral recordings, the shortest delay between the masker and test was 0.5 ms. The data revealed no noticeable difference between masking effects at ipsi- or contralateral masker source positions relative to the test source position. The amplitude-vs-delay functions for the symmetrical masker source positions looked similar, and their difference was not statistically significant for all azimuths, from 15 to 90° (from *p* = 0.17 to *p* = 0.33 as assessed by ANOVA).

**Fig 9.**
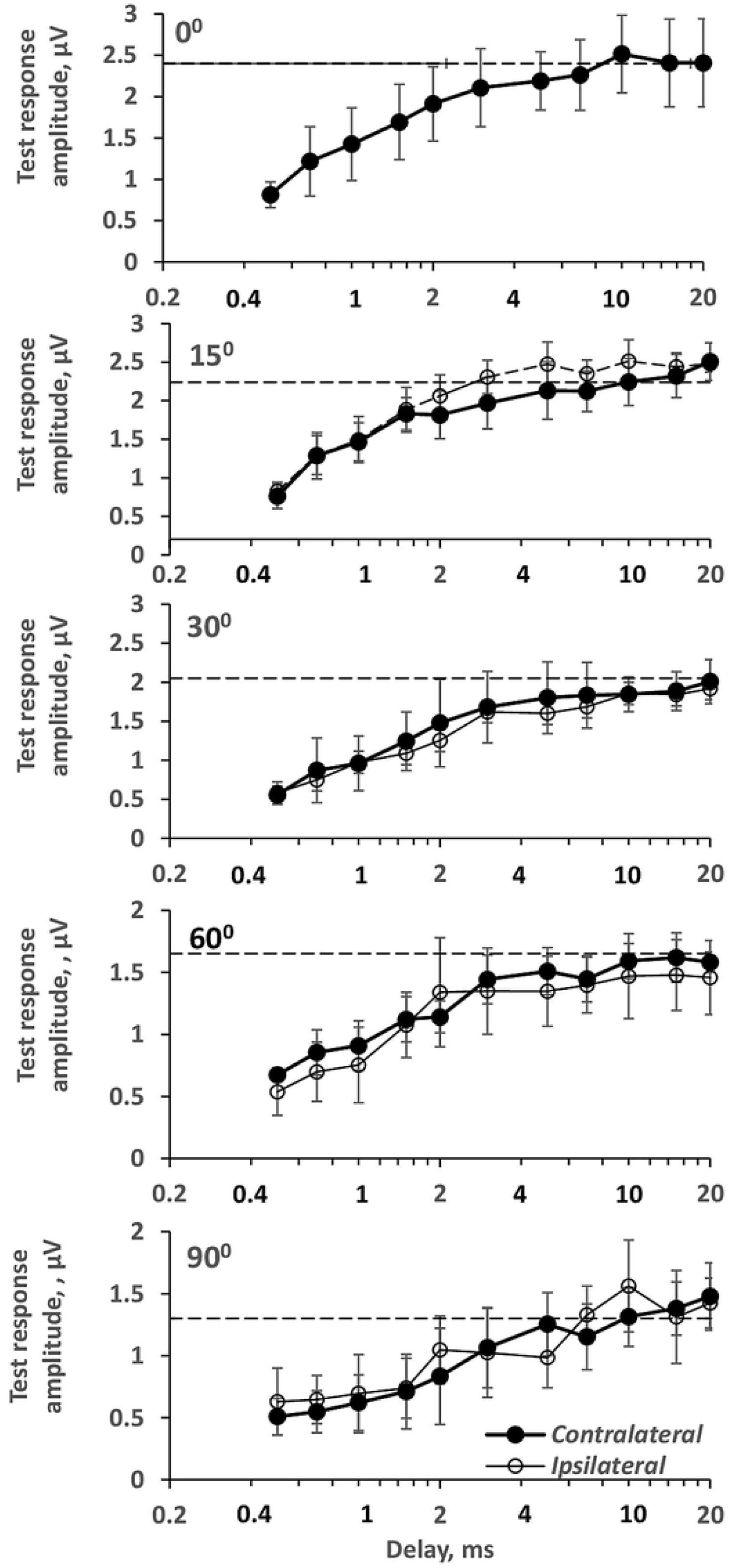
The same as Fig 7 for vertex-recorded AEPs. 0° - both masker and test stimulus sources are at the head midline.

## Discussion

In the present study, AEP release from forward masking was assessed by amplitudes of the N2.8-P3.5 component recorded from the lateral head surface and the N3.7-P4.8 component recorded from the vertex. Different latencies of these two AEP components indicate different neuronal sources of their generation. Additionally, they feature different interaural relations.

The laterally recorded N2.8-P3.5 component displayed monaural properties: it appeared in response to ipsilateral stimuli, whereas contralateral stimuli provoked components of longer latencies. Possibilities of recording a monaural AEP component from the lateral head surface in dolphins have been found earlier. Because of the shortest latency, that component was considered a response of the auditory nerve [10]. In the present study, the AEP wave that displayed properties of a response of the auditory nerve was of a small amplitude; it was not used to specify the AEP amplitude. The largest N2.8-P3.5 component might reflect the activity of peripheral auditory nuclei that have monaural inputs but are capable of generating high-amplitude evoked potentials; perhaps they are cochlear nuclei. This suggestion is supported by the manner of release from forward masking: ipsilaterally projected masker produced longer masking than the contralaterally projected masker.

The N3.7-P4.8 component was picked up from the vertex point that was symmetrical relative to both the right and left ears. Therefore, this component reflected activity provoked from any ear. Its latency was longer than that of the monaural component; this allows us to suppose its generation by higher auditory centers. Was this component a sum of activities of monaural structures excited from each ear or it reflected activity of binaurally activated auditory structures? This question may be answered by comparison of forward masking at different masker-source positions. The N3.7-P4.8 component was equally suppressed by both ipsi- and contralateral maskers. This fact may be considered evidence of converged inputs from the right and left ears with nearly equal effectiveness of both inputs for both the masker and test stimuli. As a result of equal effectiveness, forward masking might occur nearly equally when the masker and test sources were located at the same or at opposite sides relative to the head axis.

This equality may result in weaker resistance of the dolphin’s hearing to interfering sounds because sources in both the ipsi- and contralateral directions relative to the target sound produce masking. However, binaural inputs to upper auditory centers may be beneficial for the functioning of hearing during echolocation. In echolocating odontocetes, forward masking plays an important role in making the echo response invariant of the distance to the target [20]. During echolocation, forward masking of the echo response occurs because the dolphin hears its own high-level sonar pulse shortly before the low-level echo. The dependence of the echo level on distance and the dependence of the release from masking on echo delay compensate one another, thus making the echo response little dependent on distance. For targets positioned outside the head axis, the binaural inputs to auditory centers allow to keep the relation between levels of the self-heard sonar pulse and echo constant, thus providing invariance of responses of binaural auditory centers of distance.

## Acknowledgments

The study was supported by The Russian Science Foundation, Grant 22-25-00025. We thank American Journal Experts (AJE) for English language editing.

